# Efficient inference, potential, and limitations of site-specific substitution models

**DOI:** 10.1101/2020.01.18.911255

**Authors:** Vadim Puller, Pavel Sagulenko, Richard A. Neher

## Abstract

Natural selection imposes a complex filter on which variants persist in a population resulting in evolutionary patterns that vary greatly along the genome. Some sites evolve close to neutrally, while others are highly conserved, allow only specific states or only change in concert with other sites. Most commonly used evolutionary models, however, ignore much of this complexity and at best account for variation in the rate at which different sites change. Here, we present an efficient algorithm to estimate more complex models that allow for site-specific preferences and explore the accuracy at which such models can be estimated from simulated data. We find that an iterative approximate maximum likelihood scheme uses information in the data efficiently and accurately estimates site-specific preferences from large data sets with moderately diverged sequences. Ignoring site-specific preferences during estimation of branch length of phylogenetic trees – an assumption of most phylogeny software – results in substantial underestimation comparable to the error incurred when ignoring rate variation. However, the joint estimation of branch lengths, site-specific rates, and site-specific preferences can suffer from identifiability problems and is typically unable to recover the correct branch lengths. Site-specific preferences estimated from large HIV *pol* alignments show qualitative concordance with intra-host estimates of fitness costs. Analysis of site-specific HIV substitution models suggests near saturation of divergence after a few hundred years. Such saturation can explain the inability to infer deep divergence times of HIV and SIVs using molecular clock approaches and time-dependent rate estimates.

## Introduction

Over time, genome sequences change through mutations and are reshuffled by recombination. Modifications to the genomes are filtered by selection for survival such that beneficial variants spread preferentially and those that impair function are purged. As a result, some parts of genomes change rapidly, while other are strongly conserved. In addition to variation of the evolutionary rate, different sites in a genome only explore different subsets of the available states. Some positions in a protein, for example, might only allow for hydrophobic amino acids, while others require acidic side chains. Patterns of conservation and variation, possibly involving more than one site, are therefore shaped by functional constraints which in turn allows inference of biological function from genetic variation.

Similarly, phylogenetics aims at reconstructing the relationships and history of homologous sequences from the substitutions that occurred in the past. Modern phylogenetic methods describe this stochastic evolutionary process with probabilistic models of sequence evolution and aim to find phylogenies that either maximize the likelihood of observing the alignment or sample phylogenies from a posterior probability distribution (Felsenstein, 2004).

Inferring phylogenies is a computationally challenging problem since the number of phylogenies grows super-exponentially with the number of taxa and because the calculation of the likelihood is computationally costly (though linear in the number of taxa). Due to this computational complexity, the most commonly used substiution models are simple caricatures of biological complexity. The simplest substitution models assume that all sites and sequence states are equivalent and evolve at the same rate, i.e., they assume an unconstrained non-functional sequence that mutates at random between the different sequence states. Such simple models are clearly inadequate and ignoring rate heterogeneity tends to result in biased estimates of divergence times or otherwise erroneous results (Yang, 1996). Most commonly used models account for variation in substitution rates among sites and average properties of the substitution process such as transition/transversion bias or more likely substitution between similar amino acids (Nguyen *et al.*, 2015; Price *et al.*, 2010; Stamatakis, 2014; Yang, 1994). To avoid over-fitting, these methods typically don’t estimate a rate for each site, but treat site-specific rates as random effects that are integrated out (often using discrete approximations of a Gamma distribution (Yang, 1996), mixture of multiple unimodal distributions (Mayrose *et al.*, 2005), or a small number of fixed rates).

In addition to rate variation, different sites in a protein differ in amino acid they allow. Recent deep mutational scanning experiments have shown that site-specific preferences are mostly conserved between moderately diverged proteins (Doud *et al.*, 2015). Using such experimentally inferred site-specific models in phylogenetic inference greatly increases the likelihood of the data (Bloom, 2014). More than two decades ago, Halpern and Bruno (1998) pointed out that ignoring that equilibrium frequencies vary from one position to another will result in underestimation of branch lengths of a phylogeny – possibly dramatically when frequencies are heavily skewed. Hilton and Bloom (2018) recently showed that experimentally measured preference not only improve the phylogenetic fit, but also results in longer branch length estimates. Models with site-specific preferences are also known as mutation-selection balance models (Bruno, 1996; Yang and Nielsen, 2008) reflecting the intuition that equilibrium frequencies are determined by competition of diversifying processes (like mutation) and selection for an optimal function.

While biologically plausible, estimating such models from data exacerbates the over-fitting problem and it rarely attempted in practice. In the context of the site-specific models this issue has become known as *extensive parametrization* or even *infinitely many parameters problem* (Rodrigue, 2013; Spielman and Wilke, 2016). With sufficient data, however, site-specific parameters can be accurately estimated (Scheffler *et al.*, 2014; Spielman and Wilke, 2016; Tamuri *et al.*, 2012). Here, we implement an EM-style algorithm inspired by (Bruno, 1996) to infer site-specific rates and preferences from simulated data, quantify its accuracy and the different sources of bias and noise, and show how divergence time estimates depend on the fidelity with which site-specific model parameters are known. We apply this algorithm to large HIV-1 alignments and explore the consequences for phylogenetic inference.

## Efficient inference of site-specific substitution models

Following work by Halpern and Bruno (1998), we parameterize a site-specific general time-reversible (GTR) substitution model at site *a* from state *j* to *i* as:

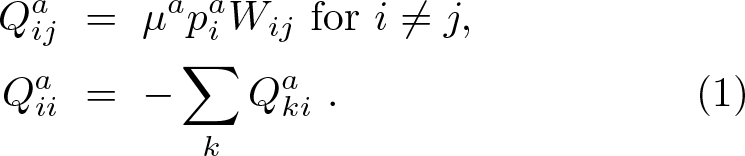

Here, *μ*^*a*^ is the substitution rate at site *a*, 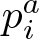 is the equilibrium frequency of state *i* at site *a*, and *W*_*ij*_ is a symmetric substitution matrix that we assume to be the same for all sites (in what follows, super script *a* will always refer to the position in the sequence, while subscript *i*, *j*, *n*, *m* refers to the state). The second equation ens es conservation of p bability. In addition, we require 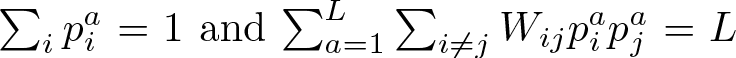 to normalize the frequencies and fix the scale of *W*_*ij*_.

Extending the approach by Bruno (1996), we show in Materials and Methods and the supplement that the iterative update rules

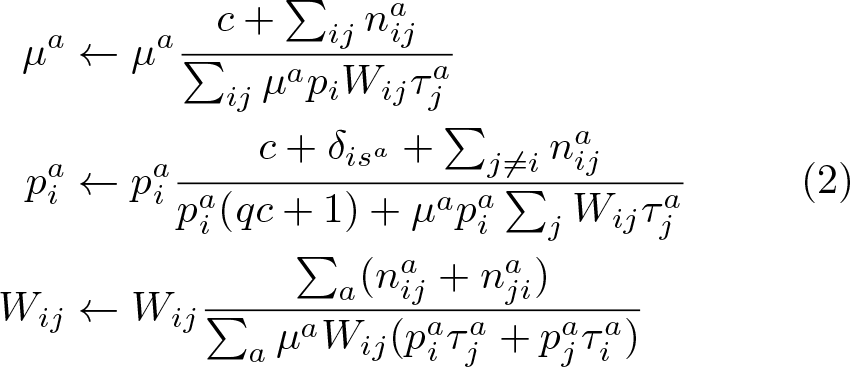

approximately maximize the likelihood of the data. Here 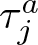 is the time site *a* spends in state *j* across the tree, 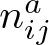 are the number of transitions from state *j* to state *i* at site *a*, *c* is a pseudocount analogous to a Dirichlet prior that in the absence of data will drive the 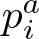 to a flat distribution and the substitution rates to *c* divided by the total tree length. To make the behavior of these update rules more explicit, we have multiplied numerator and denominator with the parameter that is being updated. This (i) shows that the update rules are multiplicative and therefore ensure positivity, and (ii) illustrates that each of these rules are the ratio of the *observed* number of transitions between states 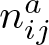 and the *expected* number 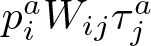 – each appropriately summed over sites or states. These update rules are an example of non-negative factorization algorithms (Lee and Seung, 2001).

## Accuracy of inferences

The validity of this iterative solution will depend (i) on the accuracy of the linear approximation, (ii) the accuracy to which 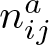 and 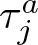 can be estimated, and (iii) the accuracy of the tree reconstruction. In addition, the Maximum Likelihood estimates might deviate from the true parameters due to over-fitting.

To assess these sources of error independently, we simulated sequences evolving along trees and explicitly recorded 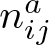 and the 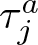 for a range of sample sizes, levels of divergence, and different degrees of variation between sites, see Materials and Methods. For each parameter set, we generated two random Yule trees, for each tree generated two random site-specific models, and for each tree/model pair two alignments, resulting in a total of eight data set per parameter combination. We quantified the accuracy of the inferences as 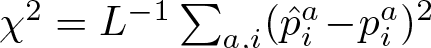 where 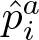 and 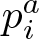 are the inferred and true equilibrium probabilities, respectively. We compared inferences using (i) the true ancestral sequences, (ii) reconstructed ancestral sequences, and (iii) reconstructions on the true phylogenetic tree.

Fig. 1 shows the average squared error χ^2^ for a trees with *n* ∈ [100, 300, 1000, 3000] taxa and a range of substitution rates as a function of the expected number of substitutions per site. The graph contains estimates of 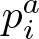 from the alignment and via the iterative scheme with correct 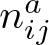 and 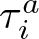 as input. The data are shown on double logarithmic scales, such that the approximately straight decrease of χ^2^ with slope −1 implies that the accuracy is inversely proportional to the expected number of substitutions. This scaling suggests that accuracy is limited by the inherent stochasticity of the evolutionary process and that the inference uses all available information efficiently. The simulation data was generated by drawing site-specific rates from a Gamma distribution with *α* = 1.5 (colored lines) and *α* = 3 (grey lines). The scaling behavior χ_2_(tree length)^−1^ is more evident for *α* = 3 than for *α* = 1.5. In case of strong rate variation, the average accuracy is dominated by a small number of sites with low rates at which frequencies are estimated poorly and the average tree length does not capture behavior at these sites. A naive approach using the frequencies of different sequences states in the alignment as estimates for 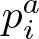 is much less accurate due to the correlation induced by shared ancestry (orange lines in Fig. 1A).

**FIG. 1.**
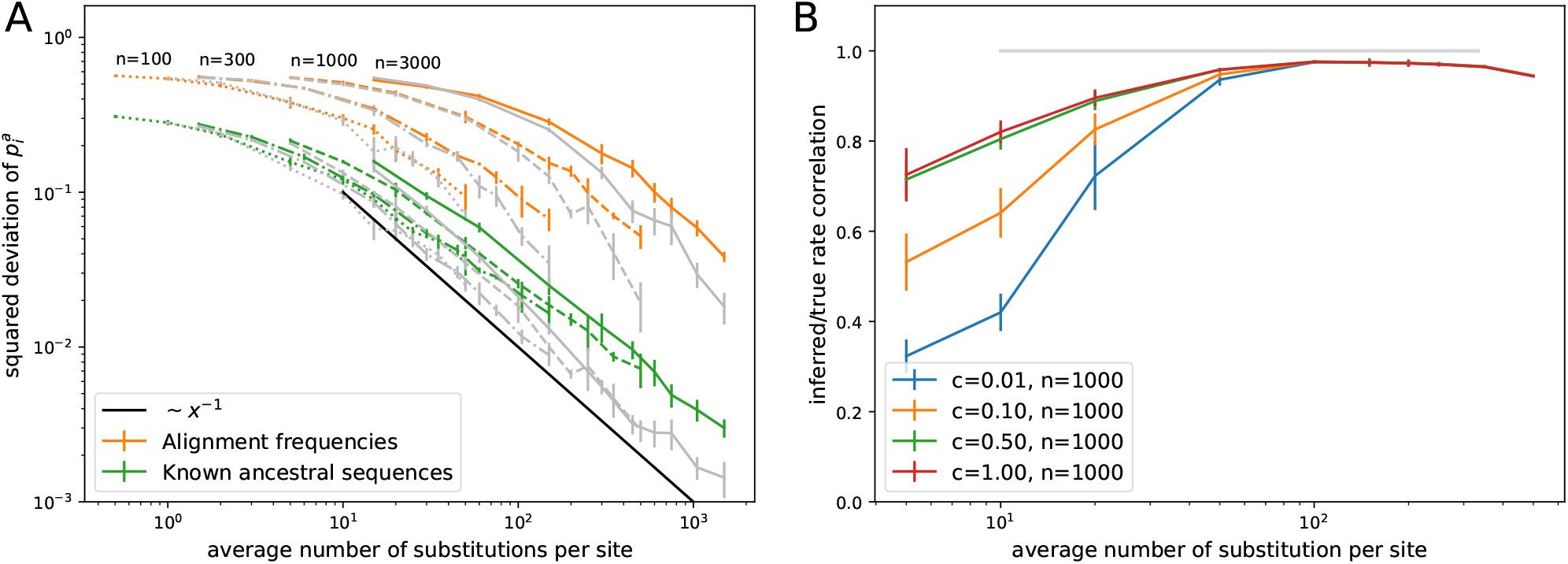
Accuracy of iterative estimation of site-specific GTR models as a function of the expected number of state changes along the tree. (A) Mean squared error of the inferred 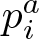 scales inversely with the tree length, suggesting the accuracy is limited by the Poisson statistics of observable mutations. Colored lines correspond to data generated assuming Gamma distributed rate variation with *α* = 1.5, gray lines with *α* = 3. The estimates are more accurate than naive estimates from the state frequencies in alignment columns. The advantage over naive estimates increases with data set size as seen from different curves with *n* ∈ [100, 300, 1000, 3000] sequences per tree. (B) The relative substitution rates are accurately inferred as soon as the typical site experiences several substitutions across the tree as quantified here as Pearson correlation coefficient between true and inferred rates. Regularization via pseudo-counts reduces over-fitting at low divergence. Analogous results for alphabets of size *q* = 20 are shown in Fig. S1.

At the largest substitution rates, branch lengths are on the order of 0.5 and the linear approximation underlying the iterative equations is not longer accurate. Nevertheless, the accuracy of the 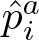 continues to increase. While equilibrium frequencies are insensitive, the substitution rate estimates are affected by linearization and Eq. 2 will consistently underestimate *μ*^*a*^. This is expected as the linearization ignores cases where the same site changes twice along a branch in very much the same way as Hamming distance will underestimate branch length. The relative substitution rates, however, are accurately estimated (with Pearson correlation coefficients > 0.9 as soon as the majority of sites experience several mutations across the tree), see Fig. 1B.

In a typical scenario, ancestral sequences are unknown and need to be inferred or summed over (marginalized). To test the influence of tree and ancestral state reconstruction, we reconstructed phylogenetic trees using IQ-tree (Nguyen *et al.*, 2015) with a GTR+R10 model (simulated nucleotide sequences) or FastTree (Price *et al.*, 2009) using the default 20 category (simulated amino-acid sequences). Fig 2 compares different schemes to reconstruct ancestral sequences, infer substitution models, and optimize the tree. The simplest approach is to take the inferred tree as given, reconstruct the ancestral states using a standard evolutionary model (e.g. Jukes-Cantor model) and calculate 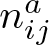 and 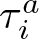 from this reconstruction. This naive approach works well up to root-to-tip distances of about 0.3, beyond which estimates deteriorate, see Fig 2 (“Reconstructed ancestral sequences”, red line). Instead of using only the most likely ancestral sequences, we can instead average over all possible ancestral states, which results in a modest improvement in accuracy (“Marginalized ancestral sequences”, purple line in Fig. 2). More significant gains are made when iterating model inference and ancestral reconstruction using the inferred site-specific model (“Iterative model estimation”, brown line). The accuracy now continues to improve up to root-to-tip distances of about one. At this level of divergence, tree reconstruction starts becoming problematic and branch lengths deviate substantially from their true values. Using the true tree instead of the reconstructed tree leads to continuous improvements of accuracy with increasing levels of divergence (yellow line). Similar improvements are achieved by optimizing tree branch lengths along with the model. We found qualitatively similar patterns for 4-letter and 20-letter alphabets in the overall accuracy of the estimated model and its dependence on the different approximations, compare Fig. 2A&B. For larger alphabets, the breakdown at large root-to-tip distances is less dramatic and sets in at larger values. However, these difference depend on the skew of the equilibrium probabilities.

**FIG. 2.**
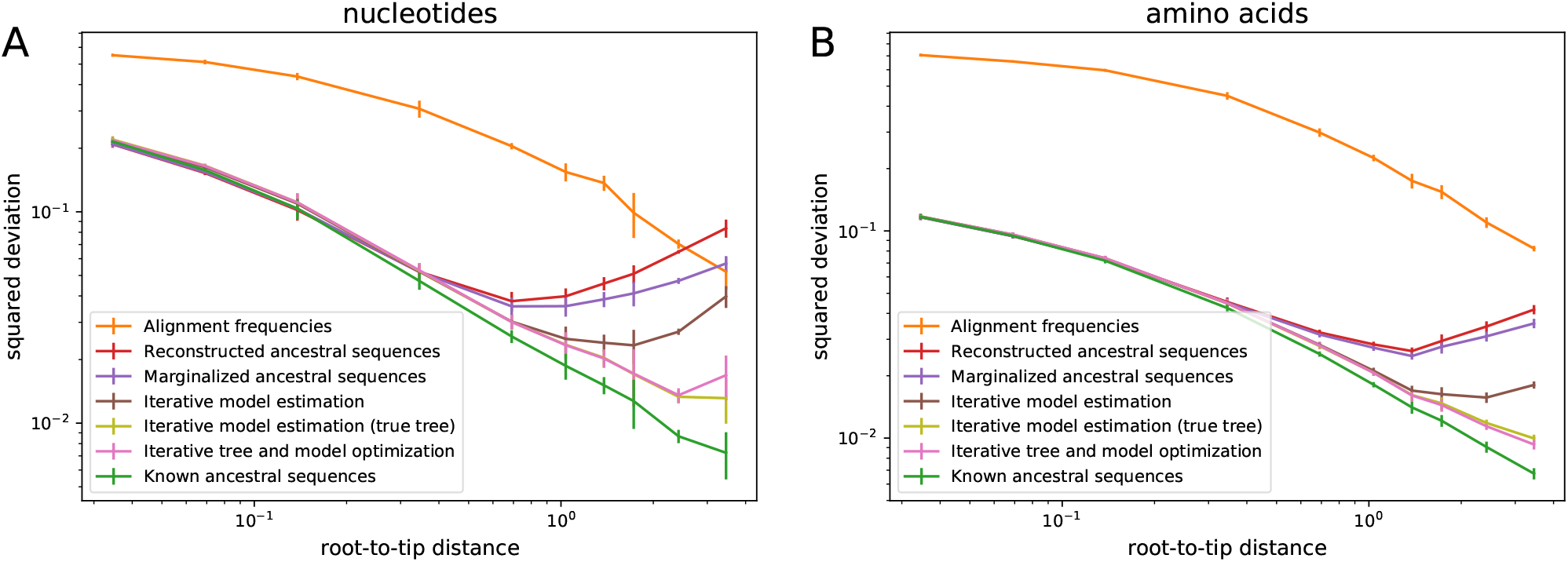
Quantification of errors stemming from tree inference and ancestral reconstruction. Panels A and B show the mean-squared deviation χ^2^ of inferred 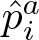 from the true 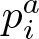 for 4-letter and 20-letter alphabets, respectively. At large root-to-tip distances, ancestral reconstruction becomes less and less certain and estimation of 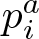 fails (red lines). These errors are gradually eliminated by first summing over ancestral uncertainty (violet), iteratively redoing ancestral reconstruction using the inferred model (brown), and re-optimizing branch lengths using the updated models (or using the true tree, yellow/pink). Data in this figure uses was generated assuming Gamma distributed rate variation with *α* = 1.5.

## Ignoring site-specific frequencies results in branch length underestimates

As previously observed by various authors (Halpern and Bruno, 1998; Hilton and Bloom, 2018), sites with heavily skewed preferences for specific states result in underestimation of branch lengths if uniform equilibrium frequencies are assumed. This is a straightforward consequence of the fact that the probability of observing the same state at random is 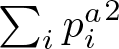 is increasing sharply with more peaked preferences. Models that don’t account for site-specific frequencies will take a large number of sites that agree between two sequences as evidence for their close evolutionary relationship, while it is simply a consequence of resampling the same states with high probability. In other words, this underestimation is the result of incorrect models of saturation and is closely related to the observation that incomplete purifying selection can result in erroneous and apparently time dependent evolutionary rates Wertheim and Kosakovsky Pond (2011).

Fig. 3 shows the average branch length (panel A) and the average root-to-tip distance of mid-point rooted trees (panel B) with branch length optimized using (i) the true model, (ii) using the GTR+R10 model of IQ-tree, and (iii) inferred models using different degrees of regularization. While branch lengths are accurately estimated when using the true model, they are systematically underestimated by the GTR+R10 model without site-specificity. Underestimation is particularly severe for the average root-to-tip distance which is dominated by long branches deep in the phylogenies which are prone to underestimation. When ignoring site-specific preferences, the inferred root-to-tip distance is essentially independent from the true value for distances greater than 1 (Fig. 3B). This effect is entirely due to skewed equilibrium frequencies, as branch length inference by IQ-tree is accurate if the simulated data had flat 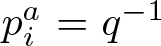 with the same rate variation. The underestimation of branch lengths is less severe for larger alphabets of if frequencies are not heavily skewed, see Fig. S2.

**FIG. 3.**
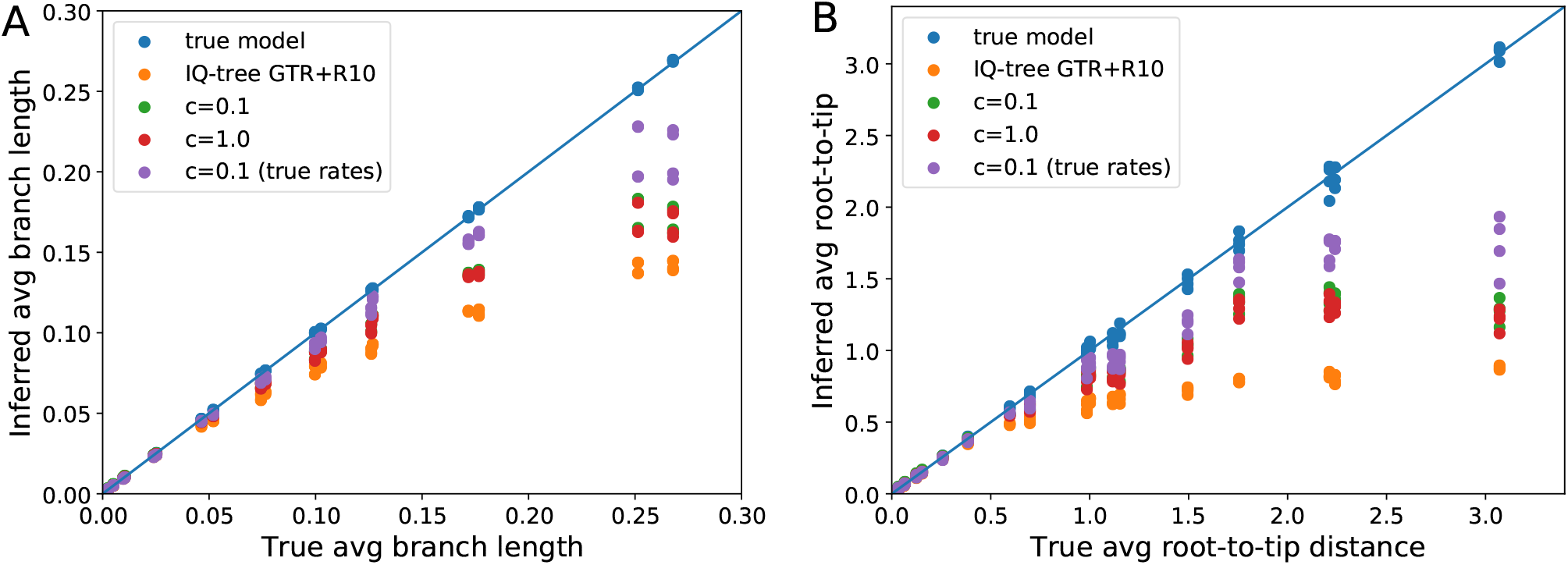
Skewed equilibrium concentration results in branch length underestimates. Panel A shows the inferred average branch length as estimated by IQ-Tree and TreeTime as a function of the true average branch length. Panel B shows the results of the same optimization for the average root-to-tip distance, which is dominated by deep long branches which are more strongly affected by underestimation. Not that inferred site-specific models only partially ameliorate underestimation, see main text. Parameters: *n* = 1000, *α* = 1.5.

Surprisingly, using the inferred site-specific models only partially rectified the problem of branch length under-estimation, despite the fact that these models are close to the true model in terms of low χ^2^. The principle contributor to this deviation are inaccuracies in the rate estimates. Combining the true rates with inferred preferences reduces the error in branch length estimation (see Fig. 3, lines labeled “(true rates)”.

## Minor model deviations can have substantial effects on divergence estimates

In the previous section, we have observed that model deviations, for example the error made during model inference, can result in substantial errors in branch length estimates. To investigate this effect more systematically, we constructed mixtures of the true model and a model with flat 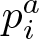 and/or *μ*^*a*^ as follows: For a mixing fraction α, we constructed a model with rates

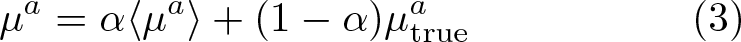

where 〈*μ*^*a*^〉 is the expectation value of the rate. Mixtures of site-specific frequencies are constructed analogously.

Fig. 4A shows how deviation from the true model affect relative branch length estimates. Deviation in rate estimates result immediately in underestimated branch length, while substantial effects of too flat site-specific preferences only manifest themselves once deviations are of order *α* = 0.3. If the same preferences are used for every site (*α* = 1), however, deviations are substantial. Deviations in rate and preferences are approximately additive. These observations are consistent with the finding above that using inferred preferences and true substitution rates result in more accurate branch length estimates than inferring both rates and preferences.

**FIG. 4.**
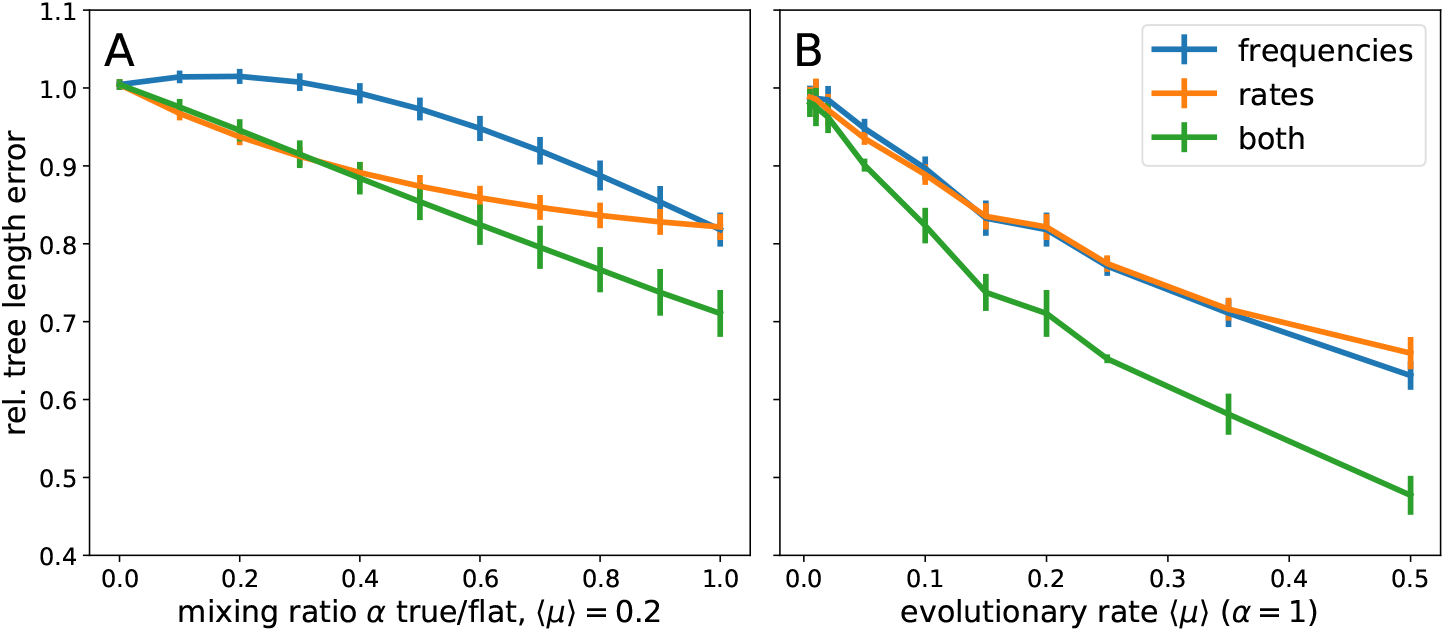
Sensitivity of branch length estimates on model misspecification. Panels A & B show the relative error in total tree length when using a mixture model as defined in Eq. 3 for branch length inference. Panel A shows this error as a function of the mixing fraction *α* for 〈*μ*〉 = 0.2. Panel B shows the error as a function of the evolutionary rate *μ* for 〈*α*〉 = 1. The mixing is applied to the equilibrium frequencies 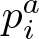, the rates *μ*^*a*^, or both. The models assume an alphabet size *q* = 4 (nucleotides).

Fig. 4B shows the degree of mis-estimation when using flat preferences, flat rates, or both as a function of the degree of divergence. The relative error increases approximately linearly with the average evolutionary rate.

## Applications to large HIV alignments

Since its zoonosis, HIV-1 has diversified into several different subtypes that differ from each other at about 20% of sites in their genome. For each subtype, thousands of sequences are available, which typically differ by about 10% (Los Alamos HIV sequence database, 2017). This large sample of moderately diverged sequences should be suited to apply our iterative inference frame work. We downloaded alignments of HIV-1 pol sequences from the LANL data base, constructed phylogenetic trees, and inferred site-specific GTR models at the nucleotide and amino acid level, see Materials and Methods.

One of the original motivations for site-specific models was the idea that equilibrium frequencies reflect the relative fitness of the different states at each position (Halpern and Bruno, 1998). Simple population genetic models predict that the fixation probability 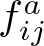 a mutation from state *j* to state *i* at site *a* should depend on the fitness difference 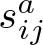 and the effective population size *N* as (Kimura, 1964)

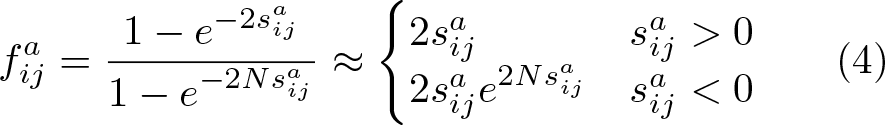

On longer time scales, the fixation rates 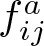 should correspond to transitions rates of the site-specific GTR model 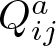. The effective population size *N* plays the role of a coalescent time scale *T_c_* and in general has little to do with census population sizes, in particular in rapidly adapting populations (Neher, 2013). Nevertheless, the logarithm of the ratio log 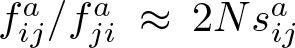 is an inter-pretable quantity related to the average fitness difference between states on time scales longer than the population genetic scale *N* ~ *T*_*c*_. We generalize this notion to multiple states and define a fitness score of state *i* at position *a* as the ratio of rates into and out of state *i*

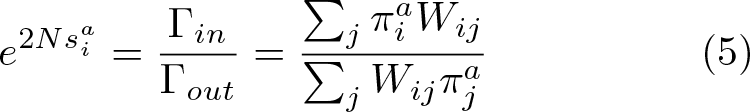

The logarithm of this ratio is expected to be proportional to the fitness difference between state *i* and the average alternative state at site *a*, while the multiplicative factor 2*N* is unknown. We compared this proxy of fitness differences to fitness costs measured using mutation selection balance models and with-in host diversity data of HIV (Zanini *et al.*, 2017). However, instead of a linear relation ship between 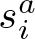 and within host estimates of fitness costs, we observed a linear relationship between the logarithm of fitness effects measured with-in host and 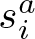 (see Fig. 5) Interestingly, this correlation between the intra-host estimates and cross-sectional estimates decreased as soon as regularization increased above 0.05 and we there for used a weak regularization of *c* = 0.01. This relationship explains about half the variance of nucleotide effects and about one third of amino acid effects.

**FIG. 5.**
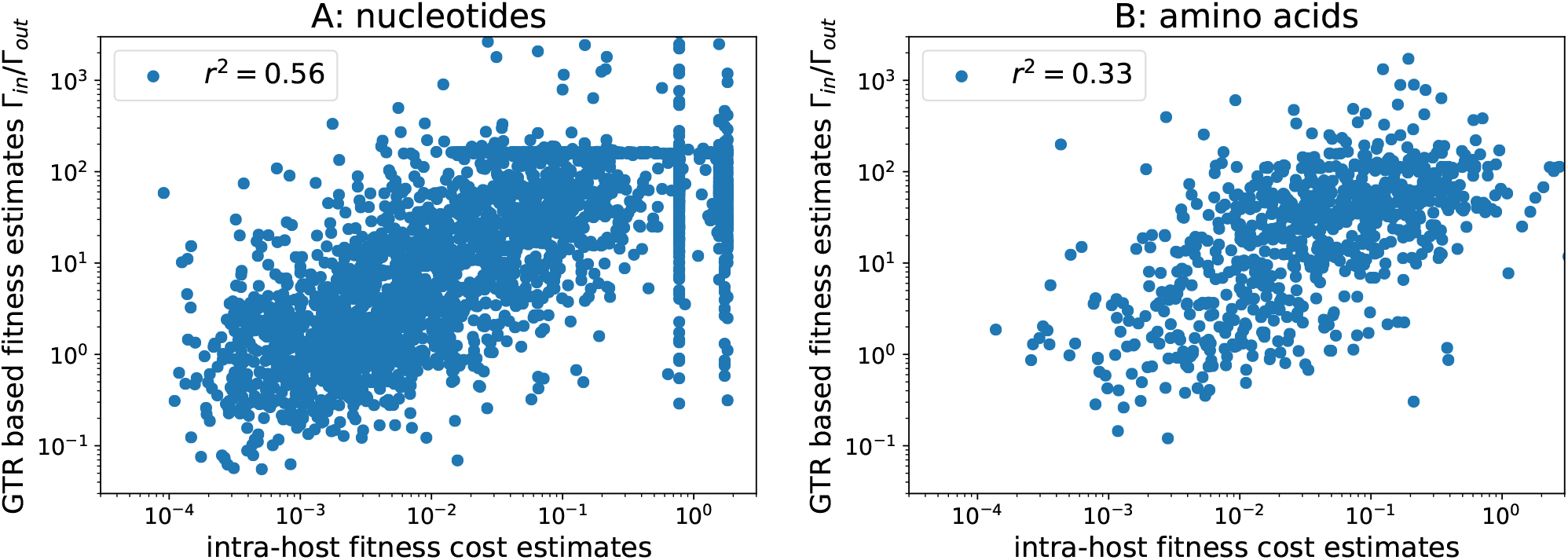
Intra-host vs cross-sectional mutation selection balance. Panel A&B shows the ratio of in/out rates for consensus nucleotides/amino acids along the *pol* of HIV-1 subtype B vs of fitness costs of non-consensus states estimated from within-host mutation selection balance. The logarithm of the rate ratio is roughly linear in the logarithm of the fitness cost. Analogous results for the genes *gag* and *nef* are shown in Fig. S3.

The fact that the relationship deviates from the expectation

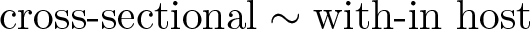

and instead is approximately

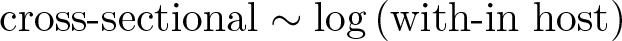

points towards different process that drive cross-sectional and with-in host diversification. While the order of fitness effects with-in host and cross-sectionally seem to be mostly concordant, with-in host variation is much greater than cross-sectional variation. Within host fitness effects are masked and damped at the population level, possibly due to fluctuating selection by diverse host immune systems or epistasis (Shekhar *et al.*, 2013; Zanini *et al.*, 2015). Such fluctuations in genetic background (epistasis) and environment (immune system) can result in non-linear averages at the population level. The ratio of in/out rates 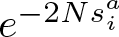 is, however, a very good correlate of alignment diversity (see Fig. S4).

Next, we quantified the effect of ignoring site-specific preferences on branch length estimates. We generated sequence pairs that evolved for a specific time *t* under the site-specific model inferred from the HIV-1 pol alignment and then estimated a maximum likelihood branch length between these two sequences. This estimate was done using a model with homogeneous equilibrium frequencies set to the average frequencies across sites while maintaining site-specific rate variation. As soon as the length of the simulated branch approaches 0.2, the length inferred by a model without site-specific preferences deviates substantially from the true value and saturates around 0.5 for larger and larger *t*. As expected, these deviations are even more severe when estimating branch length as simple sequence divergence (p-distance), while using the generating model reproduces the correct branch length.

Assuming a typical evolutionary rate of HIV of 0.002 changes per site and year, this analysis suggests that length estimates of branches longer than 100 years start to become inaccurate. Furthermore, this analysis suggests very little signal to estimate the length of branches that are longer than 300 years.

## Discussion

Sequence evolution models attempt to capture some biological realism while being computationally tractable and easy to parameterize. A common compromise is to model rate heterogeneity as random effects, while assuming identical site preferences across the sequence. Different positions in a DNA sequence or protein, however, differ markedly in the preferences for different states – this site-specific preference is the basis of all common homology search tools. While the need to account for this variation has been known for decades (Bruno, 1996; Halpern and Bruno, 1998), it is typically not modeled due to over-fitting concerns and due to computational complexity. However, amino acid preferences can now be measured with high-throughput (Fowler and Fields, 2014) and data set sizes are increasing rapidly such that more complex models can be estimated. Large data sets, however, require efficient and scalable methods.

**FIG. 6.**
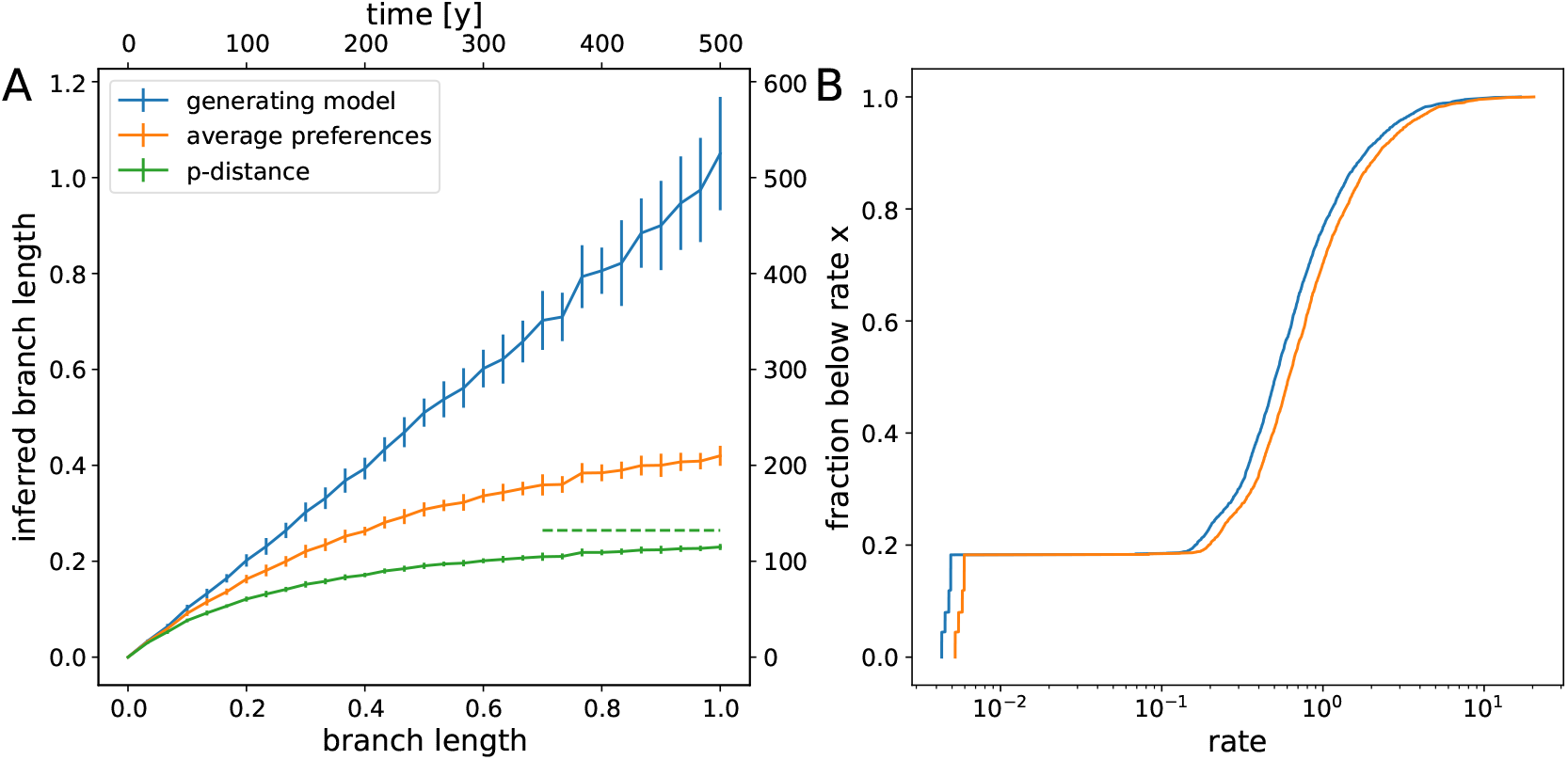
Underestimation of divergence in HIV. Panel A shows the estimated ML branch length for the model used to generate the sequences and a model with constant equilibrium frequencies as a function of true branch length. For comparison, the p-distance between the two simulated sequences is also shown. The top axis shows branch length in units of years assuming a substitution rate of 0.002/year and site. Error bars denote one standard deviation. Panel B shows the distribution of rates across sites for both models. About 20% of sites are essentially invariable, while the rates of the remainder vary by at least 10fold. The distributions differ slightly in the overall scale since rates have been rescaled such that the average substitution rate in both models is identical.

Here, we presented method to estimate site-specific GTR models using an iterative EM-type approach inspired by Bruno (1996). We showed that site-specific models can be inferred with an accuracy limited by the Poisson statistics of the substitution process using an efficient iterative scheme. For short branch lengths a parsimonious ancestral reconstruction is sufficient, while for more diverged samples iterative inference of the model, the ancestral states, and the branch lengths is necessary. The computational complexity of the inference is dominated by ancestral reconstruction and branch length optimization, which is quadratic in the alphabet size and linear in sample size and sequence length (Felsenstein, 2004). We have implemented the algorithm for site-specific model inference and branch length optimization in TreeTime (Sagulenko *et al.*, 2017).

We further explored how model choice affects the maximum likelihood estimates of branch lengths. As has been reported before, branch lengths are underestimated when variation in rate or preferences are not fully accounted for (Halpern and Bruno, 1998; Hilton and Bloom, 2018). While such underestimation often has a moderate effect on the total length of a tree, it can result in substantial underestimation of root-to-tip distance that are dominated by a few long branches close to the root. While using the correct model results in correct branch length estimates, joint inference of site-specific models and branch lengths is typically unable to recover the true branch lengths and substantial underestimation remains. This points to parameter identifiability problems rooted in the similar effects that skewed equilibrium frequencies, low rates, and shorter branch length have on the likelihood of the data.

Branch length estimates using models constructed by mixing the true model and flat Jukes-Cantor type models showed that misspecified equilibrium frequencies result in substantial errors as soon as the model deviates from the true model by 30% or more. This mirrors observations by Hilton and Bloom (2018), who found that preferences measured for influenza HA proteins of type H1 or H3 affect branch length of in the vicinity of the focal sequence, but not globally on the tree. Using site-specific models inferred from HIV alignments, we quantified the error made when ignoring site-specific preferences. While the true models are unknown in this case, this analysis nevertheless suggests that errors are substantial as soon as branch length exceed *t* = 0.2 (~100 years) and sequence divergence saturates at levels around 0.3. This result is consistent with the discrepancy between molecular clock based or biogeography based estimates of the divergence times of different SIV lineages (Worobey *et al.*, 2010): Beyond a few hundred years, there is very little signal to estimate branch length – in particular when the underlying site-specific model parameters need to be estimated themselves. This lack of information is further underscored by the average nucleotide distances between *pol* sequences of different HIV and SIV strains: HIV-1 sub-types differ at about 10% of sites, distances between pol and HIV-1 and SIVcpz are on the order of 25% and distances between sequences in the HIV-1/HIV-2/SIV compendium alignment are about 35% (all distances ignore sites with > 20% gaps). These observations are compatible with evidence for frequent reversion of HIV to a preferred sequence state following immune escape (Carlson *et al.*, 2014; Leslie *et al.*, 2004; Zanini *et al.*, 2015).

This potentially large error in branch length estimates can seriously affect deep divergence time estimates. Typically, short branches close to the tips of the tree are used to calibrate molecular clock models. The deep branches, however, tend to be much longer and rate estimation and variation of site-specific preferences will result in saturation effects not accounted for by the model (Hilton and Bloom, 2018). This effective variation in rate is related to the effect of transient deleterious mutations that inflate the rate on short time scales (Ho *et al.*, 2005; Wertheim and Kosakovsky Pond, 2011): the time it takes purifying selection to prune deleterious mutations is related to the relaxation time scales of GTR models with site-specific preferences. Instead of the *q* − 1 degenerate eigenvalues of a Jukes-Cantor model, each site has a spectrum of eigen-values and different eigenmodes relax at different speeds, generating apparently time dependent rates.

Estimating site-specific models from sequence alignments faces one fundamental problem: Reliable estimates require observation of many changes at each site which requires many sufficiently diverged sequences. At the same time, preferences at individual sites are expected to change due to epistatic interactions with other sites in the sequence, as for example witnessed by the gradual divergence of experimentally determined preferences (Doud *et al.*, 2015; Haddox *et al.*, 2018). Hence the approach described in this work is largely restricted to cases like HIV where many moderately diverged sequences (~10%) are available. Outside of this limit there either isn’t enough data to reliably estimate the large number of coefficients, or epistatic interactions need to be taken into account – probably at the expense of ignoring phylogenetic signals (Morcos *et al.*, 2011).

## Acknowledgments

We are grateful to Sarah Hilton and Pierre Barrat-Charlaix for stimulating discussions and insightful comments on the manuscript. This work was supported by core funding by the University of Basel, the Max Planck society, and ERC Stg 260686.

## Materials and Methods

### Model and notation

Most models of sequence evolution express the probability that sequence 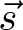 evolved from sequence 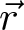 in time *t* as

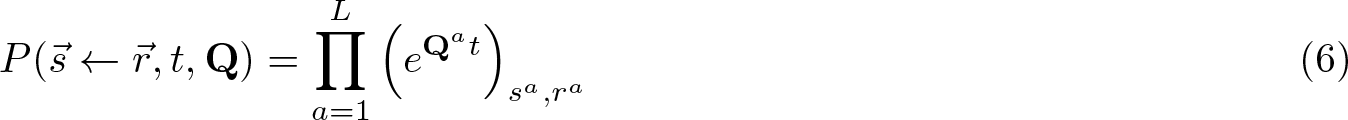

where **Q**^*a*^ is the substitution matrix governing evolution at site *a*, and *s*^*a*^ and *r*^*a*^ are the sequence states at position *a*. The product runs over all *L* sites *a* and amounts to assuming that different sites of the sequence evolve independently.

In absence of recombination, homologous sequences are related by a tree and the likelihood of observing an alignment 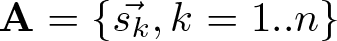 conditional on the tree *T* and the substitution model **Q**^*a*^ can be written in terms of propagators defined in Eq. 6. It is helpful to express this likelihood as product of sequence propagators defined in Eq. 6 between sequences at the ends of each branch in the tree (implicitly assuming that evolution on different branches is independent and follows the same time reversible model). Unknown sequences of internal nodes 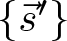 need to be summed over and the likelihood can be expressed as

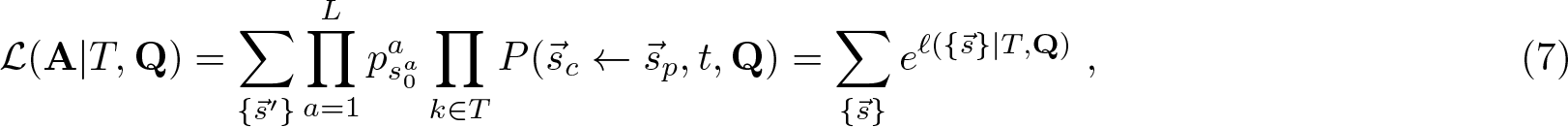

where 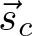 and 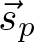 are the child and parent sequences of branch *k*, respectively, and the factor 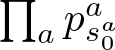 is the product of the probabilities of the root sequence 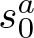 over all positions *a*. The probabilities 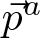 are the equilibrium probabilities of the substitution model at position *a*. The latter ensures that the likelihood is insensitive to a particular choice of the tree root. This equation defines the log-likelihood *ℓ* of a particular internal node assignment 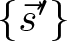 which is given by

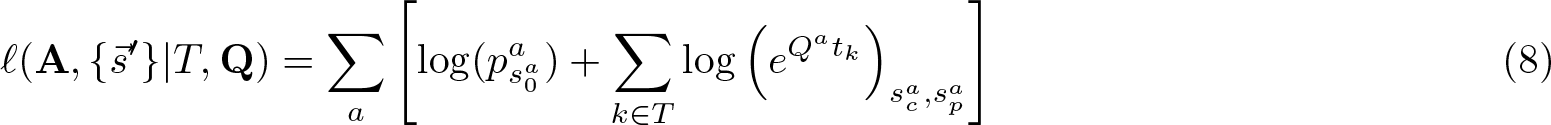

where 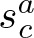 and 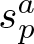 are indices corresponding to the child and parent sequence of branch *k*.

The sum over unknown ancestral sequences can be computed efficiently using standard dynamic programming techniques. Nevertheless, it requires 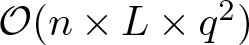 operations (where *q* is the size of the alphabet 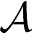) and optimizing it with respect to a large number of parameters is costly. Our goal here is to infer site-specific substitution models using a computationally efficient iterative procedure.

Instead of inferring completely independent models for every site in the genome, we follow Halpern and Bruno (1998) and only allow for site-specific rates and equilibrium frequencies while using the same transition matrix for every site. Such a site-specific general time-reversible (GTR)) model, can be parameterized as:

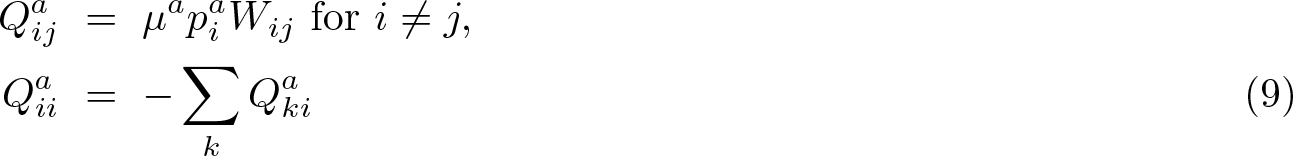

where *W*_*ij*_ is a symmetric matrix with *W*_*ii*_ = 0 and the second equation ensures conservation of probability. In addition, we require 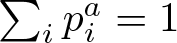 and 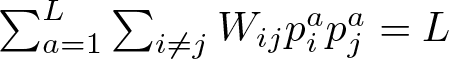 to ensure that the average rate per site is *μ*^*a*^.

### Iterative optimization algorithm

The derivatives of 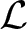 with respect to *μ*^*a*^, 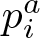, and *W*_*ij*_ need to vanish at the values that maximize 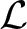.

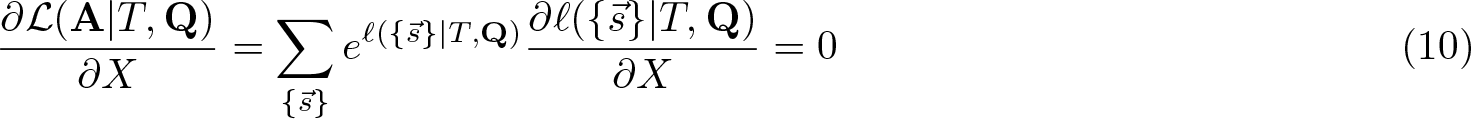

where *X* is one of the parameters we vary.

These conditions can be solved iteratively. Here, we derive the update for *μ*^*a*^ and refer to the supplement for the other update rules. The derivative of the log-likelihood for a specific sequence assignment is given by

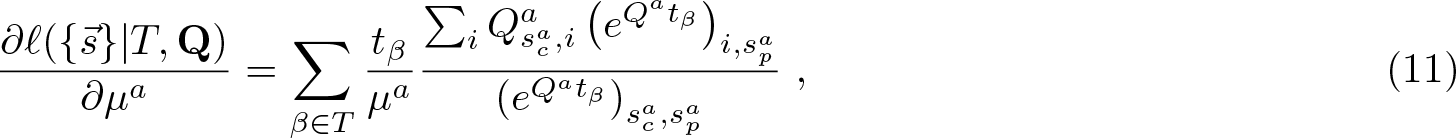

where 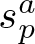 and 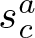 are the states at site *a* at the parent and child end of branch *β*. The individual terms in this sum behave very differently for cases where 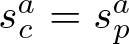 (no change at site *a* on branch *β*) and 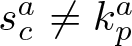 (at least one mutation). In the limit of short branches *μ*^*a*^*t*_*β*_ ≪ 1, we can expand the matrix exponential *e*^**Q***t*^ = *δ*_*ij*_ + **Q***t* + … to obtain approximate but solvable conditions for maximum likelihood parameter estimates, see supplement. We will separate the sum over branches into those with 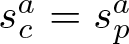 and 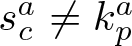. Suppressing the index *a* for the position in the sequence, we find

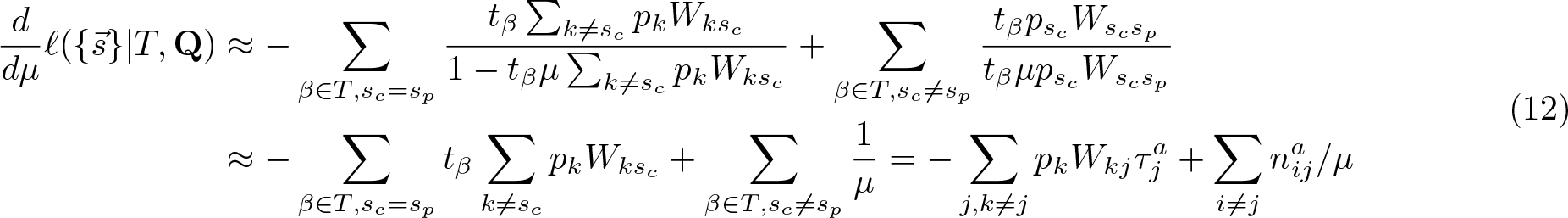

where 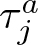 is the sum of all branch length along which site *a* is in state *j* and 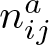 is the number of times the sequence at site *a* changes from *j* to *i* along branches of the tree (we have re-instantiated the position index *a* in the last line). Additional terms necessary for regularization and normalization are discussed in the supplement. Setting this using expression to zero (and the corresponding ones in the supplement) suggests solving for *μ*^*a*^ at fixed 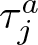 and 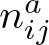 using the iterative update rules given in Eqs. 2. The quantities 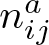 and 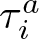 can be averaged over unknown ancestral states.

### Implementation

We extended our package TreeTime (Sagulenko *et al.*, 2017) to handle site-specific GTR models by adding an additional class `GTR_site_specific` that generalizes the `GTR` class. Since these classes have an almost identical interface and can be used interchangeably in other analysis run by TreeTime. Using the new class, TreeTime can generate sequences with site-specific evolutionary models and infer these models from sequence data using the algorithms above.

### Generation of simulated data

To generate models and sequence ensembles for which the ground truth is known, we sampled binary trees with *n* = 100, 300, 1000, 3000 leaves and Yule tree branch length statistics using betatree (Neher *et al.*, 2013) for values of the average substitution rate given by ⟨*μ*⟩ = [0.005, 0.01, 0.02, 0.05, 0.1, 0.15, 0.2, 0.25, 0.35, 0.5].

For each tree, we sampled two site-specific models for sequences of length *L* = 1000 from following distribution: Site-specific rates *μ*^*a*^ were sampled iid from a Gamma distribution with parameter *α* = 1.5 or 3.0. Site-specific preferences 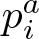 (or equilibrium probabilities) were sampled from Dirichlet distributions with parameters *α* = 1 for alphabets with *q* = 4 states (nucleotides) or *α* = 0.2 and *α* = 0.5 for alphabets with *q* = 20 (amino acids). The entries of the transition matrix *W*_*ij*_ were sampled from a Dirichlet distribution with *α* = 2. The average substitution rate of these models was fixed to the required ⟨*μ*⟩. In addition, we generated one set of models (*q* = 4) with rate variation but uniform 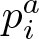 and *W*_*ij*_. For each combination of tree and model, two sets of sequences were evolved using the sequence generation function of TreeTime.

In total, this amounts to 80 alignments for each of the data set sizes *n* = 100, 300, 1000, 3000 and four different ensembles. For each alignment, we reconstructed phylogenetic trees using IQ-tree (Nguyen *et al.*, 2015) with a GTR+R10 model or FastTree (Price *et al.*, 2009) using the default 20 category model for nucleotide and amino-acid sequences, respectively. These trees were used to infer site-specific models from data, see below. The exact workflow is documented in the script `src/generate_toy_data.py` in the associated git repository at github.org/neherlab/2019_Puller_SiteSpecificGTR.

### Model inference from simulated data

Simulated data and trees (reconstructed or true) were read in by TreeTime and models reconstructed using functions of TreeTime to infer models with the different approximations discussed in the text. The exact workflow is documented in the script `src/reconstruct_toy_data.py` in the associated git repository at github.org/neherlab/2019_Puller_SiteSpecificGTR.

### HIV sequence analysis

HIV-1 sequences were downloaded from LANL HIV database (Los Alamos HIV sequence database, 2017) setting filters to “one sequence per patient”, “non-ACGT<0.3”. Separate downloads where made for sequences the following sets: *pol*, subtype B (2019-05-05); *pol*, subtype C (2019-05-13); *gag*, *nef*, subtype B (2019-06-07).

Sequences were aligned to the HXB2 reference sequences using mafft (Katoh and Standley, 2013), phylogenies were inferred using IQ-tree (Nguyen *et al.*, 2015), and ancestral sequences were inferred using TreeTime (Sagulenko *et al.*, 2017) via the nextstrain’s augur pipeline (Hadfield *et al.*, 2018). The different steps were assembled into a pipeline using the workflow manager Snakemake (Köster and Rahmann, 2012).

The sequences and trees were then used for site-specific GTR inference as implemented in TreeTime (Sagulenko *et al.*, 2017). The scripts detailing this analysis and producing the figures are available on GitHub in repository github.org/neherlab/2019_Puller_SiteSpecificGTR.

