## Supplementary text and figures for "Efficient inference, potential, and limitations of site-specific substitution models"

(Dated: January 18, 2020)

For a given set of sequences $\{\vec{s_k}\}$ consisting of the alignment at the tips and the ancestral sequences, the log-likelihood is given by

$$\ell(\{\vec{s}\}|T, \mathbf{Q}) = \sum_a \left[\log(p^a_{s^a_0}) + \sum_{\beta\in T} \log\left(e^{Q^a t_\beta}\right)_{s^a_c, s^a_p}\right] \tag{1}$$

where $\mathbf{Q}$ is the site specific substitution model consisting of rates, frequencies, and a symmetric transition matrix

$$\begin{aligned} Q^a_{ij} &= \mu^a p^a_i W_{ij} \text{ for } i \neq j, \\ Q^a_{ii} &= -\sum_k Q^a_{ki} \, . \end{aligned} \tag{2}$$

At the maximum, the derivatives of $\ell$ with respect to each of the model parameters should vanish when averaged over all internal site assignments. To facilitate the algebra below, we summarize the short branch length assumption and the associated linear approximations here:

$$\left(e^{Qt}\right)_{ij} = \delta_{i,j} + \sum_{m=1}^{\infty} \frac{t^m}{m!} \left(Q^m\right)_{ij} \approx \begin{cases} 1 - t\mu^a \sum_{k\neq i} p^a_k W_{ki} & i = j \\ t\mu^a p^a_i W_{ik} & i \neq j \end{cases} \tag{3}$$

The derivatives of $\left(e^{Qt}\right)_{ij}$ with respect to the different model parameters are therefore

$$\frac{d}{d\mu^a}\left(e^{Q^a t}\right)_{ij} \approx \delta_{ab} \begin{cases} -t \sum_{k\neq i} p^a_k W_{ki} & i = j \\ t p^a_i W_{ik} & i \neq j \end{cases} \tag{4}$$

$$\frac{d}{dp^a_n}\left(e^{Q^a t}\right)_{ij} \approx \delta_{ab} \begin{cases} -t\mu^a(1-\delta_{i,n}) W_{ni} & i = j \\ t\mu^a \delta_{i,n} W_{nk} & i \neq j \end{cases} \tag{5}$$

$$\frac{d}{dW_{mn}}\left(e^{Q^a t}\right)_{ij} \approx \begin{cases} -\delta_{n,j}(1-\delta_{m,i}) t\mu^a p^a_m & i = j \\ \delta_{m,i}\delta_{n,j} t\mu^a p^a_m & i \neq j \end{cases} \tag{6}$$

Since the matrix $W_{ij}$ is symmetric, the derivative with respect to $W_{ij}$ and $W_{ji}$ are the same. We will account for this symmetry below explicitly and refrain from carrying the symmetric term through lengthy derivations. Note that the derivatives with respect to $W_{ii}$ vanish as required.

##### Update rule for rates $\mu^a$

Only site $a$ will contribute to the derivative of $\ell$ with respect to $\mu^a$. We will therefore drop all indices referring to the position and re-instantiate the index later. As discussed in the main text, we will separate the sum over branches $\beta$ into those where sequence is the same of parent and child ($s_c = s_p$) and those where the sequence differs ($s_c \neq s_p$).

$$\begin{aligned} \frac{d}{d\mu}\ell(\{\vec{s}\}|T, \mathbf{Q}) &\approx -\sum_{\beta\in T, s_c = s_p} \frac{t_\beta \sum_{k\neq s_c} p_k W_{ks_c}}{1 - t_\beta \mu \sum_{k\neq s_c} p_k W_{ks_c}} + \sum_{\beta\in T, s_c\neq s_p} \frac{t_\beta p_{s_c} W_{s_c s_p}}{t_\beta \mu p_{s_c} W_{s_c s_p}} \\ &\approx -\sum_{\beta\in T, s_c = s_p} t_\beta \sum_{k\neq s_c} p_k W_{ks_c} + \sum_{\beta\in T, s_c\neq s_p} \frac{1}{\mu} = -\sum_{j, k\neq j} p_k W_{kj}\tau_j + \sum_{i\neq j} n_{ij}/\mu \end{aligned} \tag{7}$$

Here, $\tau_j$ is the sum of all branch lengths on which $s_c = j$ and $n_{ij}$ is the number of times the sequence changes from $j$ to $i$ somewhere along the tree. Setting this expression to 0 and re-instantiating the site index $a$, we get

$$\mu^a = \frac{\sum_{i\neq j} n_{ij}^a}{\sum_{j,k\neq j} p_k^a W_{kj} \tau_j^a} \tag{8}$$

##### Update rule for frequencies $p_i^a$

As above, only site $a$ will contribute to the derivative of $\ell$ with respect to $p_n^a$. We will therefore drop all indices referring to the position and re-instantiate the index later.

$$\begin{aligned}
\frac{d}{dp_n}\ell(\{\vec{s}\}|T,\mathbf{Q}) &\approx \frac{\delta_{s_0,n}}{p_n} - \sum_{\beta\in T, s_c=s_p} \frac{t_\beta \mu (1-\delta_{n,s_c}) W_{ns_c}}{1 - t_\beta \mu \sum_{k\neq s_c} p_k W_{ks_c}} + \sum_{\beta \in T, s_c \neq s_p} \delta_{n,s_c} \frac{t_\beta \mu W_{ns_p}}{t_\beta \mu p_{s_c} W_{s_c s_p}} \\
&\approx \frac{\delta_{s_0,n}}{p_n} - \sum_{\beta\in T, s_c=s_p} t_\beta \mu (1-\delta_{s_c,n}) W_{s_c n} + \sum_{\beta\in T, s_c=n, s_c\neq s_p} \frac{1}{p_n} \\
&= \frac{\delta_{s_0,n}}{p_n} - \sum_{j\neq n} \tau_j \mu W_{jn} + \sum_{j\neq n} \frac{n_{nj}}{p_n}
\end{aligned} \tag{9}$$

Reinstantiating the position index $a$, we find

$$p_n^a = \frac{\delta_{n,s_0^a} + \sum_j n_{nj}^a}{\sum_{j\neq n} \mu W_{nj} \tau_j^a} \tag{10}$$

##### Update rule for the transition matrix $W_{ij}$

Since $W_{ij}$ does not depend on the position, the derivative of $\ell$ with respect to $W_{ij}$ depends on all sites in the alignment.

$$\begin{aligned}
\frac{d}{dW_{mn}}\ell(\{\vec{s}\}|T,\mathbf{Q}) &\approx -\sum_a \sum_{\beta\in T, s_c^a=s_p^a} \frac{\delta_{n,s_p^a}(1-\delta_{m,s_c^a}) t_\beta \mu^a p_m^a}{1 - t_\beta \mu \sum_{k\neq s_c^a} p_k W_{ks_c^a}} + \sum_{\beta\in T, s_c\neq s_p} \delta_{m,s_c} \delta_{n,s_p} \frac{t_\beta \mu^a p_m^a}{t_\beta \mu^a p_m^a W_{mn}} \\
&\approx -\sum_a \sum_{\beta\in T, s_c^a=s_p^a} \delta_{n,s_p^a}(1-\delta_{m,s_c^a}) t_\beta \mu^a p_m^a + \frac{1}{W_{mn}} \sum_a n_{mn}^a \\
&= \sum_a \tau_n^a \mu^a p_m^a + \frac{1}{W_{mn}} \sum_a n_{mn}^a
\end{aligned} \tag{11}$$

Identifying $W_{nm}$ and $W_{mn}$ then results in the following symmetric expression

$$\frac{d}{dW_{mn}}\ell(\{\vec{s}\}|T,\mathbf{Q}) \approx \sum_a \mu^a(\tau_n^a p_m^a + \tau_m^a p_n^a) + \frac{1}{W_{mn}} \sum_a (n_{mn}^a + n_{nm}^a) \tag{12}$$

Setting this expression to 0 and solving for $W_{ij}$ we find

$$W_{mn} = \frac{\sum_a n_{mn}^a + n_{nm}^a}{\sum_a \mu^a(\tau_n^a p_m^a + \tau_m^a p_n^a)} \tag{13}$$

##### Implementation of the constraint $\sum_i p_i^a = 1$

Equilibrium frequencies need to be normalized to one and constraints of this nature can be accounted for by Lagrange parameters $\lambda^a$ coupling to the constraint:

$$\frac{d}{dp_n}\left[\ell(\{\vec{s}\}|T,\mathbf{Q}) + \lambda^a \sum_i p_i^a\right] = 0 \tag{14}$$

The constraint results in a modified extremal condition

$$0 = \delta_{s_0^a,n} - p_n \sum_{j\neq n} \tau_j^a \mu^a W_{jn} + \sum_{j\neq n} n_{nj}^a + p_n^a \lambda^a \tag{15}$$

Summing this condition over $n$, we find

$$0 = 1 - \sum_{i,j;i\neq n} \mu^a \tau_j^a p_i^a W_{ji} + \sum_{i,j;i\neq j} n_{ij}^a + \lambda^a \tag{16}$$

At the extremum of $\mu^a$, we have $\sum_{i,j;i\neq n} \mu^a \tau_j^a p_i^a W_{ji} = \sum_{i,j;i\neq j} n_{ij}$ and therefore $\lambda^a = -1$. With the constraint $\sum_i p_i^a = 1$, the update rule therefore becomes

$$p_n^a = \frac{\delta_{n,s_0^a} + \sum_j n_{nj}^a}{1 + \sum_{j\neq n} \mu W_{nj} \tau_j^a} \tag{17}$$

In practice it is often helpful to explicitly enforce normalization after each iteration.

#### Regularization

When there are very few mutations at a particular site we need to regularize the problem to avoid overfitting and numerical instabilities. To this end, we assume a Dirichlet prior of the form

$$\prod_i (p_i^a)^c \tag{18}$$

which drives frequency estimates away from the boundary for $c > 0$. The derivative of the logarithm of this prior is $c/p_i^a$ and adding this term to the update rule results in

$$p_n^a = \frac{\delta_{n,s_0^a} + c + \sum_j n_{nj}^a}{1 + qc + \sum_{j\neq n} \mu W_{nj} \tau_j^a} \tag{19}$$

where $q$ is the size of the alphabet. In an analogous fashion, we add a weak regularization to the update rules of $\mu^a$ and $W_{ij}$.

#### Summation over unobserved states

So far, we have derived conditions that maximize the likelihood as a function of parameters of the substitution model for a fixed assignment of sequences at internal nodes. In practice, the internal nodes are unknown and can either be fixed to the most-likely state or summed over. Since the approximate maximum likelihood conditions are linear, this average over unknown ancestral states simply amounts to replacing $\tau_i^a$ and $n_{ij}^a$ by their averages which can be efficiently computed by standard recursive algorithms.

### Supplementary figures

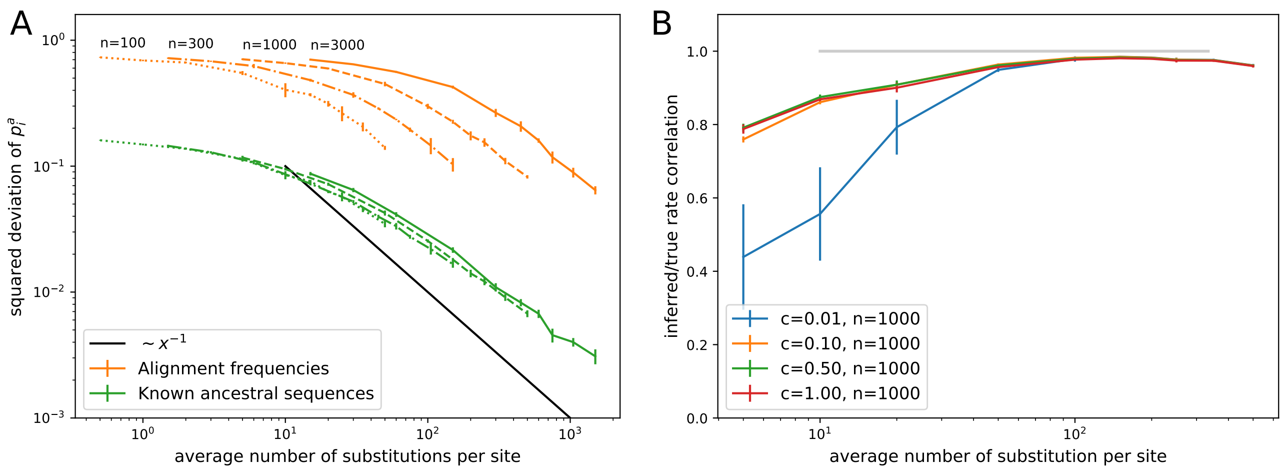


FIG. S1 Accuracy of iterative estimation as a function of the total tree length, i.e., the expected number of state changes along the tree, for amino acid alphabets. (A) Mean squared error of the inferred $p_i^a$ scales inversely with the tree length, suggesting the accuracy is limited by the number of observable mutations. The estimates are vastly more accurate than naive estimates from the state frequencies in alignment columns. Different curves are for $n \in [100, 300, 1000]$ sequences. (B) The relative substitution rates are accurately inferred as soon as the typical site experiences several substitutions across the tree as quantified here as Pearson correlation coefficient between true and inferred rates. Regularization via pseudo-counts reduces over-fitting at low divergence.

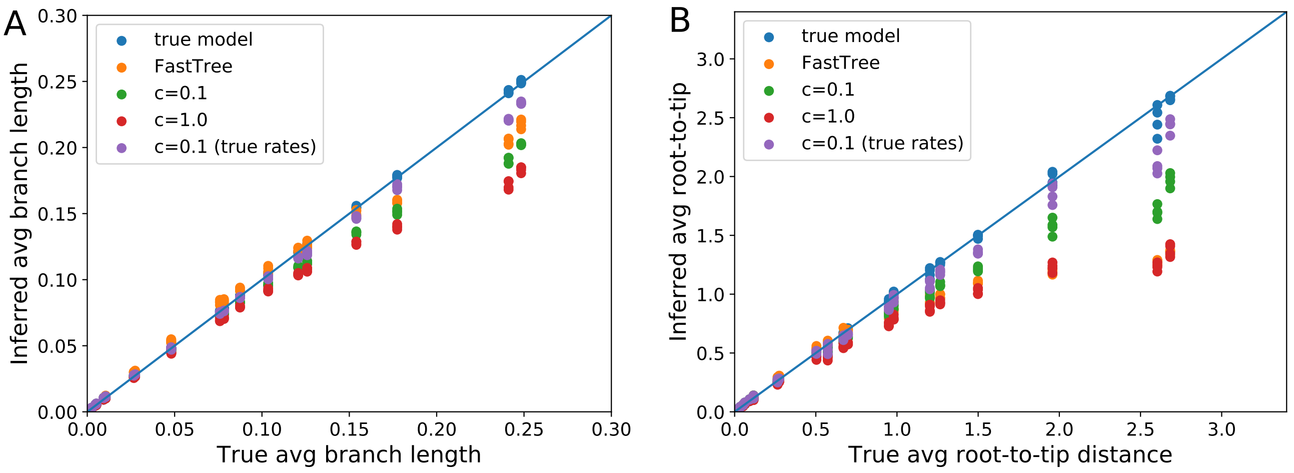


FIG. S2 **Skewed equilibrium concentration results in branch length under estimates – amino acid alphabet**. Panel A shows the inferred average branch length as estimated by IQ-Tree and TreeTime as a function of the true average branch length. Panel B shows the results of the same optimization for the average root-to-tip distance, which is dominated by deep long branches which are more strongly affected by underestimation. Note that inferred site-specific models only partially ameliorate underestimation, see main text.

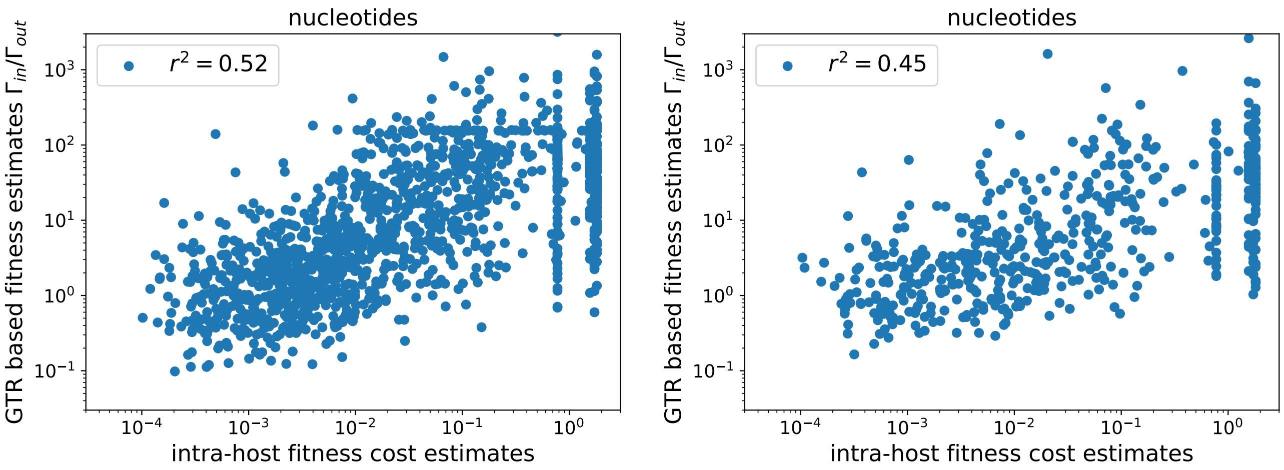


FIG. S3 **Intra-host vs cross-sectional mutation selection balance.** In analogy to Fig. 5, the panels A&B show the ratio of in/out rates for consensus nucleotides acids along the *gag* and *nef* gene of HIV-1 subtype B vs of fitness costs of non-consensus states estimated from within-host mutation selection balance. The logarithm of the rate ratio is roughly linear in the logarithm of the fitness cost.

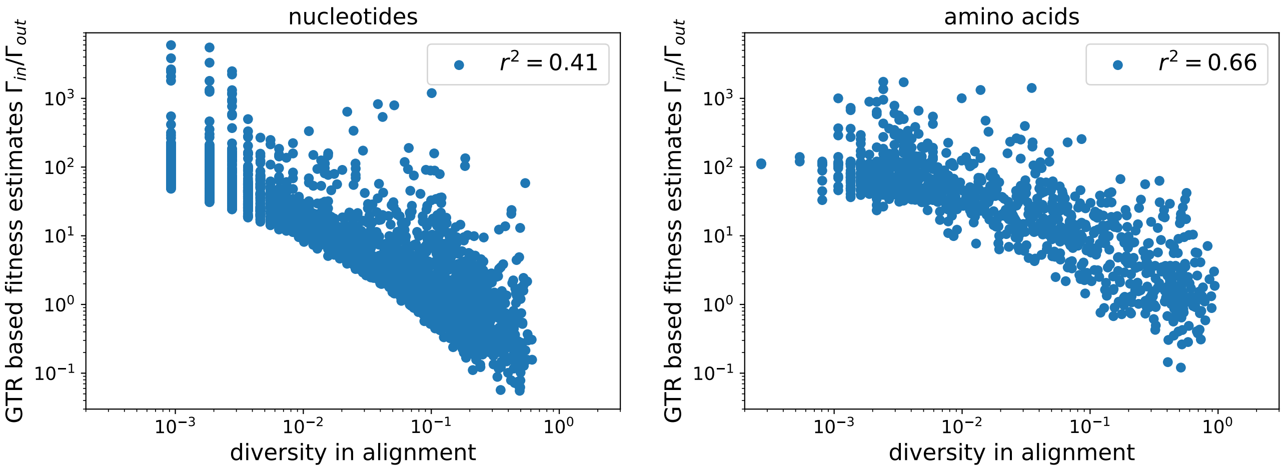


FIG. S4 **Diversity and GTR rate estimates.** Panel A&B shows the ratio of in/out rates for consensus nucleotides/amino acids along the *pol* of HIV-1 subtype B vs the diversity in columns of the corresponding alignment.
